# Phenological mismatch between trees and wildflowers: Reconciling divergent findings in two recent analyses

**DOI:** 10.1101/2023.08.01.551551

**Authors:** Benjamin R. Lee, Evelyn F. Alecrim, Jessica R.K. Forrest, J. Mason Heberling, Richard B. Primack, Risa D. Sargent

## Abstract

1. Recent evidence suggests that community science and herbarium datasets yield similar estimates of species’ phenological sensitivities to temperature. Despite this, two recent studies by Alecrim et al. (2023) and Miller et al. (2022) found contradictory results when investigating an identical ecological mechanism (phenological mismatch of wildflower flowering and of shading by deciduous trees; “phenological escape”) with separate datasets.
2. Here, we investigated whether differences between the two studies’ results could be reconciled by testing four hypotheses related to model design, species selection, spatiotemporal data extent, and phenophase selection.
3. Hybrid model structures brought results from the two datasets closer together but did not fully reconcile the differences between the studies. Cropping the datasets to match spatial and temporal extents appeared to reconcile most differences but only at the cost of much higher uncertainty associated with reduced sample size. Neither species selection nor phenophase selection seemed to be responsible for differences in results.
4. *Synthesis:* Our analysis suggests that although species-level estimates of phenological sensitivity may be similar between crowd-sourced and herbarium datasets, inherent differences in the types and extent of data may lead to contradictory inference about complex biotic interactions. We conclude that, until community science data repositories grow to match the range of climate conditions present in herbarium collections or until herbarium collections grow to match the spatial extent and temporal frequency of community science repositories, ecological studies should ideally be evaluated using both datasets to test the possibility of biased results from either.

## INTRODUCTION

A trend of earlier flowering and leafing-out of plants in the temperate zone of the Northern Hemisphere in response to warmer spring temperatures has emerged as one of the most sensitive and well-documented biological indicators of climate change (Chmielewski & Rötzer, 2001; Inouye, 2022; Parmesan, 2007). This shift in spring phenology has wide-ranging implications for ecosystem services and processes (Kim et al., 2018; Piao et al., 2019; Richardson et al., 2013) as well as for ecological interactions involving pollinators, insect herbivores, and seed dispersers (e.g., Freimuth et al., 2022; Ren et al., 2020; Simmonds et al., 2020). Species may react differently to warming, potentially altering phenological relationships to the detriment of one or more of the species, a situation sometimes termed “phenological mismatch” (Renner & Zohner, 2018). Ecologists are actively investigating this phenomenon and discussing what constitutes proof of its existence (e.g., Iler et al., 2021; Kharouba & Wolkovich, 2023; Samplonius et al., 2021). Phenological mismatch is challenging to measure, and evidence for it is limited, with most studies focusing on mismatches between trophic levels (i.e., “trophic mismatch”), especially with respect to plant-pollinator interactions (Renner & Zohner, 2018).

Recently, a paper by Heberling et al. (2019), using data gathered as far back as the 1850s, concluded that trees and spring wildflowers in eastern North America are increasingly exhibiting a phenological mismatch because tree phenology is more responsive to spring temperature compared to that of spring-blooming wildflowers. Spring-blooming wildflowers rely on access to early seasonal light to assimilate more than half—and up to 100%—of their annual carbon budgets (Lapointe, 2001); phenological mismatch could disrupt access to this critical resource via unequal changes in tree and wildflower phenology in response to changing climate conditions. One of the key findings of the Heberling et al. study is that wildflowers are experiencing less spring light than they did in the past because of lower phenological responsiveness to spring temperature in comparison with trees. Further, this window of spring sunlight is expected to become even shorter in coming decades, with projected reductions in energy budgets, survival, and reproductive success of the wildflower species (Heberling et al., 2019; Lee et al., 2022). Because species in the herbaceous understory make up most of the plant diversity in North American forests (Gilliam, 2007; Spicer et al., 2020, 2022), a projected decline in energy budget could have serious consequences for the conservation of biodiversity in these forests as well.

While the Heberling et al. (2019) study was innovative, one limitation was that all the phenological observations were from a single location: the town of Concord, Massachusetts, USA. Would the pattern of phenological mismatch between trees and wildflowers shown for this one place be shared across larger geographical ranges encompassing wider variation in environmental conditions? Recently, two articles written by two independent research teams investigated this very question across eastern North America (Alecrim et al., 2023; Miller et al., 2022). Surprisingly, the two papers reached opposing conclusions (Fig. 1A). The paper by Miller et al. (2022), using data from herbarium specimens collected between 1870 and 2019, concluded that eastern North American trees were more responsive to a warming climate than were spring-blooming herbs, similar to the single-site results of the Heberling et al. (2019) study. In contrast, the study by Alecrim et al. (2023), using phenological data collected by community scientists (USA National Phenology Network; https://www.usanpn.org/usa-national-phenology-network) between 2009 and 2021, found that understory wildflowers were more responsive to a warming climate than were trees—suggesting that wildflowers would actually experience a longer window of spring sunlight as temperatures rise. While data from community science projects and herbaria can be correlated and reveal similar patterns (e.g., Iwanycki Ahlstrand et al., 2022; Ramirez-Parada et al., 2022; Spellman & Mulder, 2016), the articles by Miller et al. (2022) and Alecrim et al. (2023) suggest that these different data sources may sometimes lead to starkly different conclusions when applied to complex ecological interactions.

**Fig. 1.**
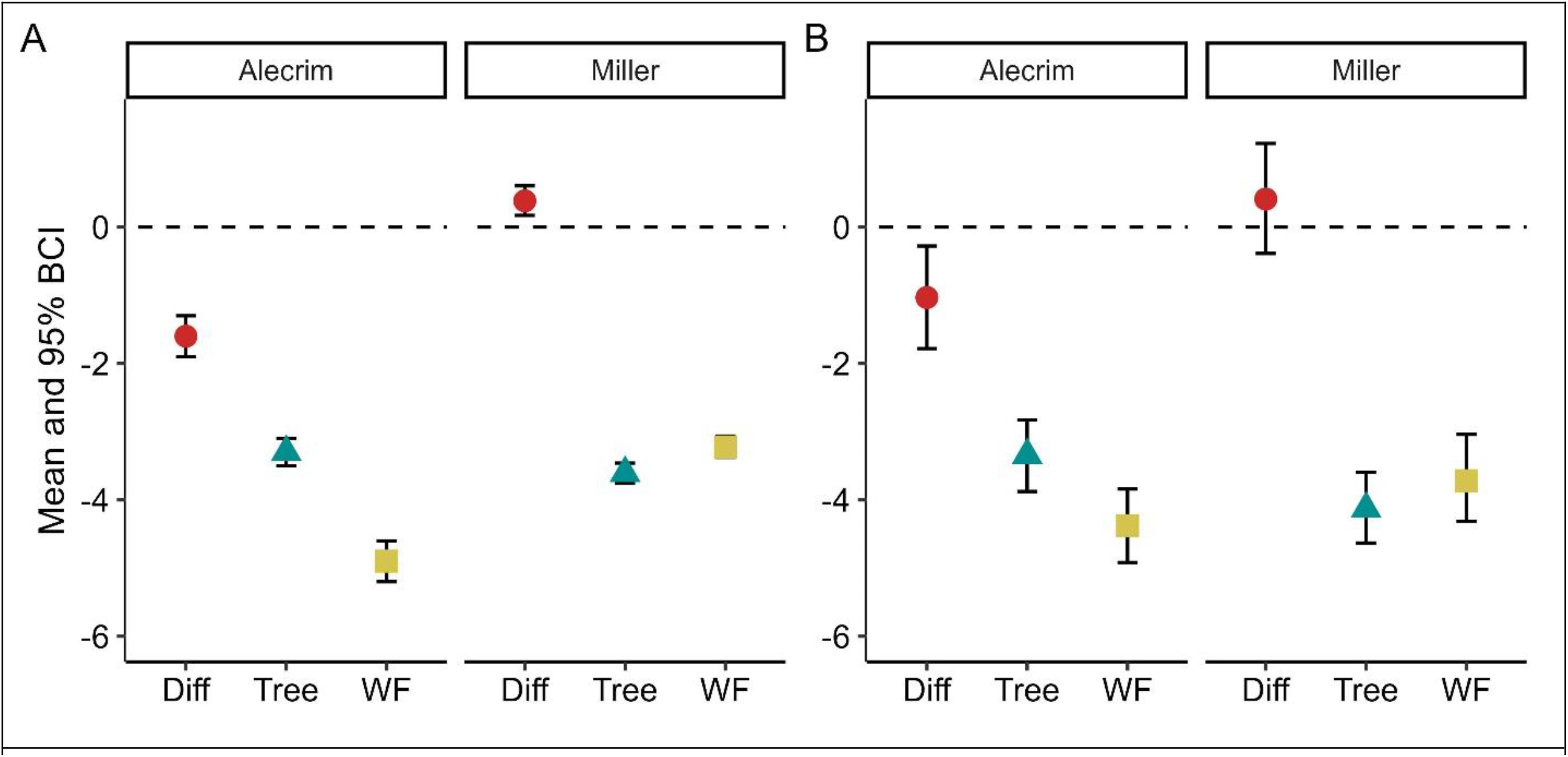
Posterior estimated means (points) and 95% Bayesian credible intervals (whiskers) of phenological sensitivities of wildflowers (WF; yellow squares), trees (green triangles), and the difference between them (red circles), (A) as reported by Alecrim et al. (2023) and Miller et al. (2022), and (B) controlling for temperature window (Mar–Apr), historical temperature data estimates (PRISM), model structure (*DOY*_*[i]*_ *=β*_*s[i]*_ *× temperature*_*[i]*_ + α_*s*_ + α_*k*_), and statistical program. A negative difference indicates that herbs have higher sensitivity to spring temperature than trees; a positive difference indicates the reverse. Posterior parameter estimates for latitudinal/temperature bins for both the original and merged model structures are provided in Fig. S1.

For the present Forum article, many of the authors of these two papers, and of a related paper (Lee et al., 2022), came together to discuss what factors (summarized in Table 1) may have contributed to the contradictory results and to test whether controlling for some of these factors could lead to better agreement between the two papers’ analyses and conclusions. Ultimately, we aimed, if possible, to reconcile the contrasting results from these two studies, which would also help to inform the use and interpretation of herbarium and community science data in future phenological studies.

**Table 1.**
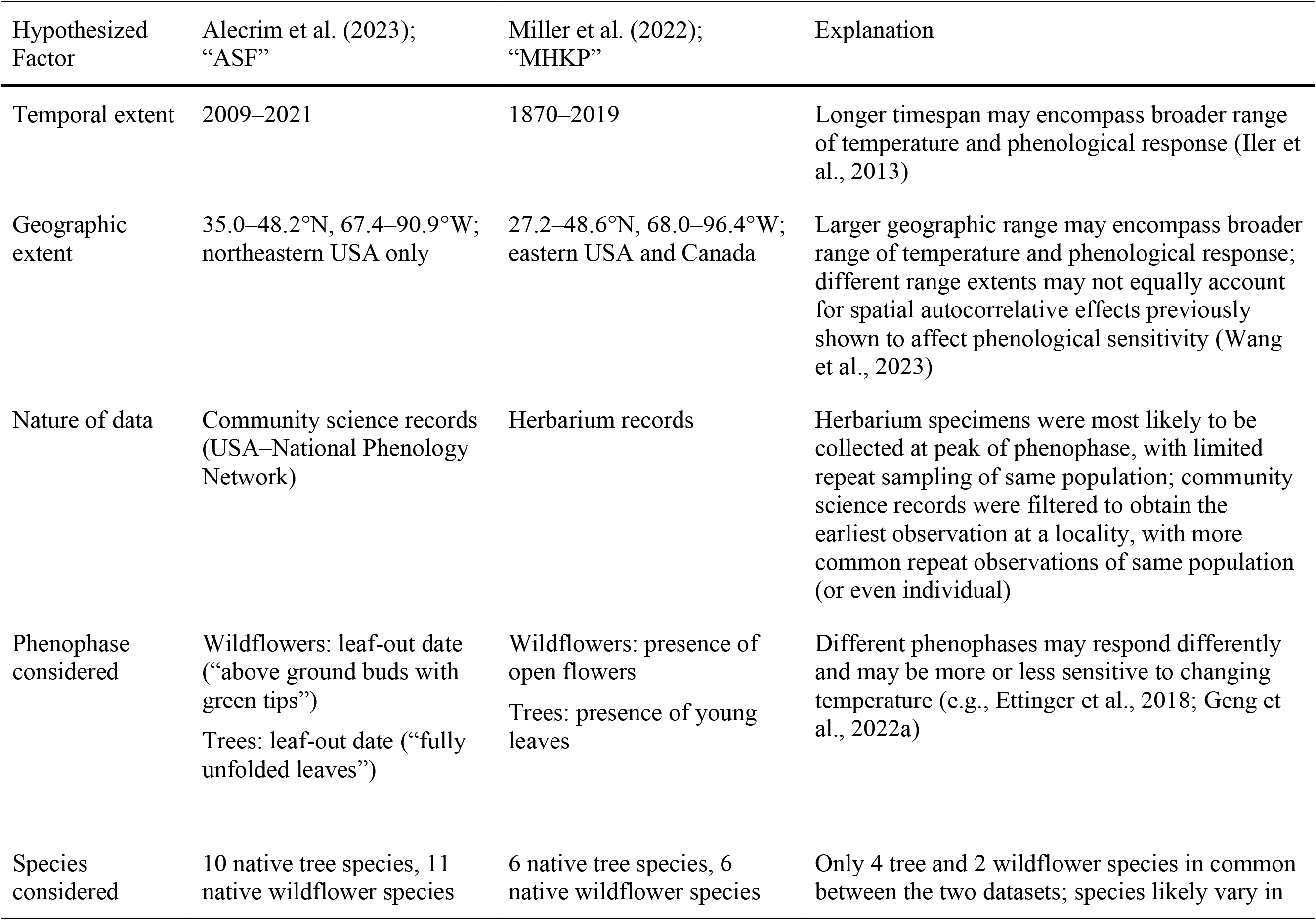
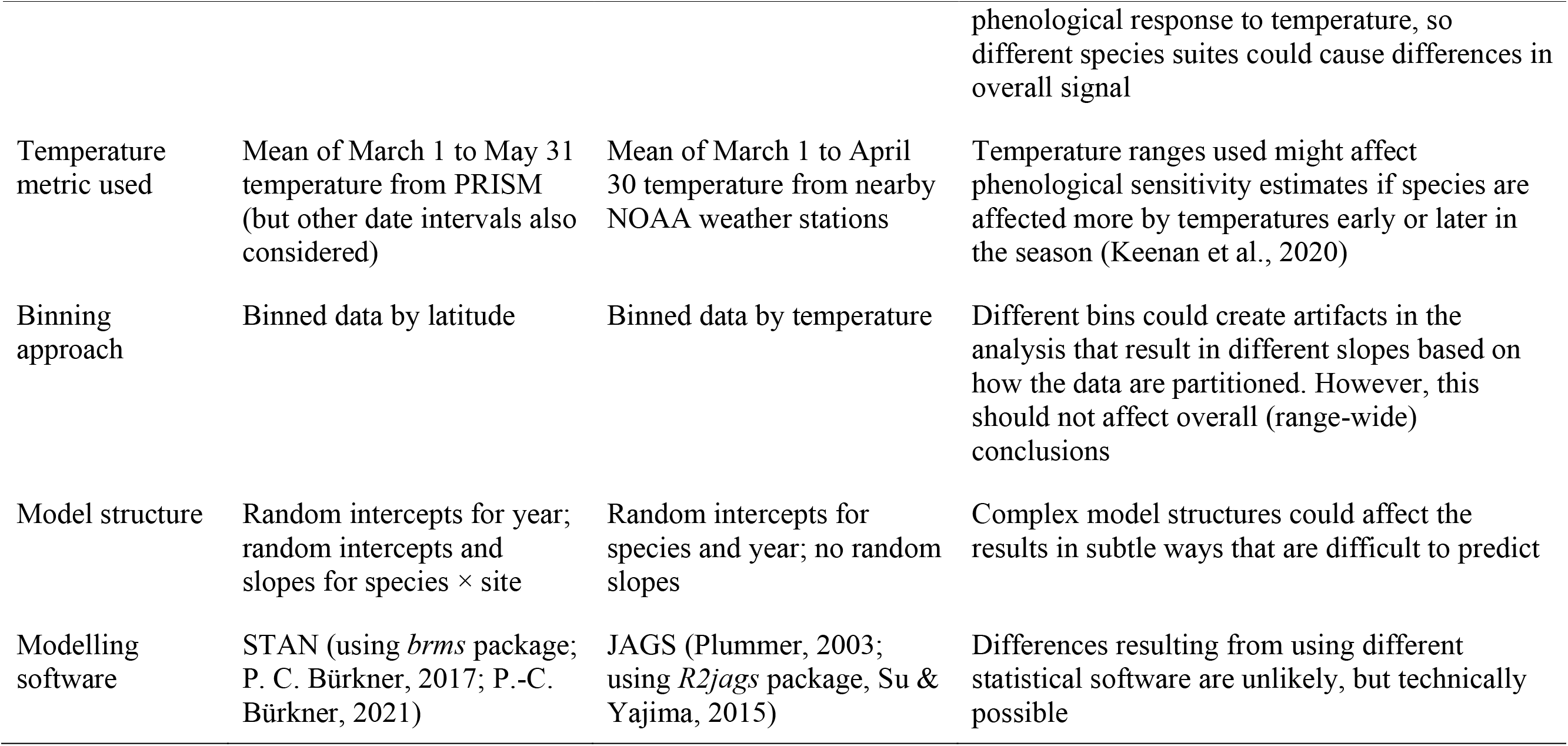
Differences between Alecrim et al. (2023) and Miller et al. (2022) studies.

| Hypothesized Factor | Alecrim et al. (2023); “ASF” | Miller et al. (2022); “MHKP” | Explanation |
| --- | --- | --- | --- |
| Temporal extent | 2009–2021 | 1870–2019 | Longer timespan may encompass broader range of temperature and phenological response (Iler et al., 2013) |
| Geographic extent | 35.0–48.2°N, 67.4–90.9°W; northeastern USA only | 27.2–48.6°N, 68.0–96.4°W; eastern USA and Canada | Larger geographic range may encompass broader range of temperature and phenological response; different range extents may not equally account for spatial autocorrelative effects previously shown to affect phenological sensitivity (Wang et al., 2023) |
| Nature of data | Community science records (USA–National Phenology Network) | Herbarium records | Herbarium specimens were most likely to be collected at peak of phenophase, with limited repeat sampling of same population; community science records were filtered to obtain the earliest observation at a locality, with more common repeat observations of same population (or even individual) |
| Phenophase considered | Wildflowers: leaf-out date (“above ground buds with green tips”)<br>Trees: leaf-out date (“fully unfolded leaves”) | Wildflowers: presence of open flowers<br>Trees: presence of young leaves | Different phenophases may respond differently and may be more or less sensitive to changing temperature (e.g., Ettinger et al., 2018; Geng et al., 2022a) |
| Species considered | 10 native tree species, 11 native wildflower species | 6 native tree species, 6 native wildflower species | Only 4 tree and 2 wildflower species in common between the two datasets; species likely vary in phenological response to temperature, so different species suites could cause differences in overall signal |
| Temperature metric used | Mean of March 1 to May 31 temperature from PRISM (but other date intervals also considered) | Mean of March 1 to April 30 temperature from nearby NOAA weather stations | Temperature ranges used might affect phenological sensitivity estimates if species are affected more by temperatures early or later in the season (Keenan et al., 2020) |
| Binning approach | Binned data by latitude | Binned data by temperature | Different bins could create artifacts in the analysis that result in different slopes based on how the data are partitioned. However, this should not affect overall (range-wide) conclusions |
| Model structure | Random intercepts for year; random intercepts and slopes for species × site | Random intercepts for species and year; no random slopes | Complex model structures could affect the results in subtle ways that are difficult to predict |
| Modelling software | STAN (using *brms* package; P. C. Bürkner, 2017; P.-C. Bürkner, 2021) | JAGS (Plummer, 2003; using *R2jags* package, Su & Yajima, 2015) | Differences resulting from using different statistical software are unlikely, but technically possible |

In our comparison of these two studies, we first identified several hypotheses to explain the incongruencies in results: (1) differences in the model structure, environmental condition estimates, and modelling programs used; (2) differences in the species from which data were available; (3) differences in spatiotemporal extents of the data used (and consequent differences in the temperature ranges considered); and (4) differences in the phenophase measured (i.e., flowering versus first emergence of foliage of the wildflowers). Here, we address these points using a comparative modelling approach and quantitative analyses. We then discuss the implications of our findings for other studies that use long-term data to assess phenological mismatch.

## METHODS AND ANALYSIS

### H1. Differences in model approaches

Studies of how climate change affects phenological mismatch typically compare the reaction norms (phenological sensitivities) of interacting species to the same climatic driver (often temperature). Indeed, both Miller et al. (2022; hereafter MHKP) and Alecrim et al. (2023; hereafter ASF) used similar statistical approaches to quantify the sensitivity of spring wildflower and canopy tree phenology to average spring temperature, which is known to be a major driver of plant phenology in spring in eastern North America (Flynn & Wolkovich, 2018; Lee & Ibáñez, 2021b, 2021a; Polgar et al., 2014; Sevenello et al., 2020) However, there were some nuanced differences in model structure. Specifically, ASF’s primary model structure estimated responses of Day of Year (DOY) to spring temperature (spanning March–May) with hierarchical group interactions (*j*; species × site level; equivalent to random effects in mixed modelling approaches) shaping estimates of model slope and intercept. ASF’s analysis also added a year random effect (*k*) to the intercept to account for non-independence of observations from the same year:

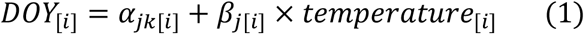

In contrast, MHKP used a different metric of spring temperature (spanning March–April). Further, these authors used a common intercept (*β*_*0*_) for all species within each functional group (i.e., spring-blooming wildflowers vs. trees), did not include random effects in the slope estimates, and included random intercepts only for species (*s*) and year (*k*), but no interaction with site since herbarium collections and measurements are not typically repeated at the site level:

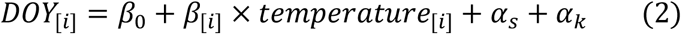

These differences in model structure and in the definition of spring temperature could have contributed to the observed differences in overall estimates of phenological sensitivity (Table 1).

Importantly, both groups of authors ran binned analyses aimed at quantifying differences in phenological sensitivity at southern, mid, and northern latitudes. ASF separated observations by latitude (prioritizing spatial congruency), while MHKP instead binned observations based on their associated March–April mean temperatures (prioritizing environmental similarity). These different binning approaches could lead to between-study differences in trends observed within latitudinal or climatic zones; however, they would not on their own explain why ASF found range-wide greater temperature sensitivity in wildflowers while MHKP found greater temperature sensitivity in trees. For this reason, we primarily focus this article on differences in range-wide sensitivities. Still, we did find some nuanced differences associated with binning strategy and we have included bin-specific graphs in the Supplemental Materials. These differences are also summarized briefly in the H3 section below.

Lastly, while both analyses used similar hierarchical Bayesian modeling approaches, MHKP conducted their analysis using JAGS software, which approximates the posterior distribution using a Gibbs sampler, whereas ASF used STAN, which approximates the posterior distribution using a Hamilton Monte Carlo sampler. Previous studies suggest it is unlikely that the different programs and samplers would yield substantially different results (Monnahan et al., 2017), but it is still possible that they could be responsible for the discrepancies in posterior estimates noted above and thus should be controlled for.

To test the influence of these differences in model structure, programming environment, and spring temperature window, we reran the analyses using a single common spring temperature window (here we used average March–April temperatures extracted from the PRISM database), and statistical program (STAN). We used a hybrid model structure that primarily reflects the MHKP model structure (Eq. 2), but with the addition of random slope effects for species:

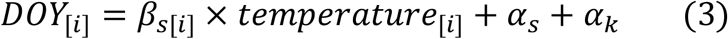

### H1 results

When controlling for model structure, programming environment, and spring temperature window, the sensitivities (i.e., phenological reaction norms) of spring-blooming wildflowers and trees become more similar for the ASF dataset (Fig. 1B). However, the results still indicate that spring-blooming wildflowers have greater temperature sensitivity than trees (95% BCI: −1.78 to −0.28; does not overlap zero). Conversely, for the MHKP dataset, the difference between functional groups becomes non-significant (95% BCI: −0.39 to 1.23; overlaps zero). These findings indicate that the different modelling approaches or temperature calculations explain some, but not all, of the differences between the two original studies.

### H2. Differences in species investigated

Phenological sensitivity has been shown to vary systematically across different groups of plants based on shared traits, growth strategies, and genetics (Panchen et al., 2014; Rafferty & Nabity, 2017). And while phenological sensitivity is commonly aggregated by groups of species (as with the two papers analyzed here), the most common level at which phenological sensitivity is reported is at the species level. This is important because sensitivity can vary substantially across species within the same functional group. For example, although Alecrim et al. (2023) found an average phenological sensitivity of −4.9 days °C^*-1*^, species-specific sensitivities ranged between −5.4 and −3.0 days °C^*-1*^, suggesting large differences in outcomes with respect to changes in spring light availability. Some species like *Sanguinaria canadensis* were extremely sensitive (and therefore more likely to gain access to light) while others, such as *Trillium grandiflorum*, were less sensitive and thus more likely to experience reductions in access to spring light.

ASF and MHKP each derived the overall sensitivity of wildflower and tree phenology by averaging across different groups of species in these functional groups. ASF collated observations for more species (11 wildflower species and 10 tree species) than did MHKP (six of each group). Four tree species were included in both studies (*Acer rubrum, Acer saccharum, Fagus grandifolia*, and *Quercus rubra*), but only two wildflower species were common between them (*Erythronium americanum* and *Sanguinaria canadensis*). Because MHKP did not include species-level random slopes, we are unable to compare species-level sensitivities from the original models. Instead, we compare the species-level reaction norms for the hybrid binned model design (Eq. 3) for the wildflowers and trees in each study (Fig. 2).

**Fig. 2.**
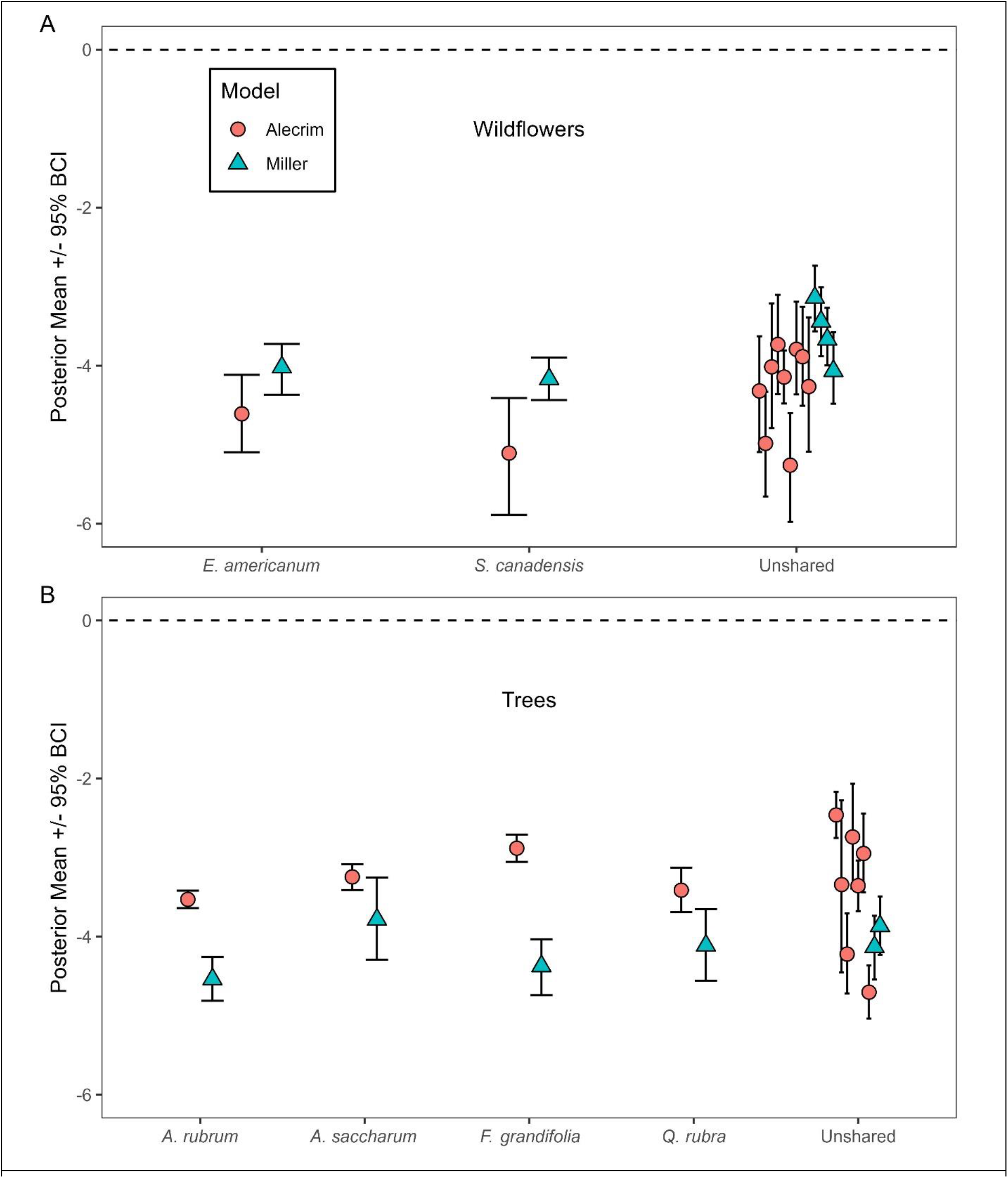
Species-level reaction norms (in days °C^*-1*^) for (A) wildflower species and (B) tree species in the Alecrim et al. (2023; red circles) and Miller et al. (2023; blue triangles) datasets. Named species were included in both papers’ species pools while all species unique to each paper are grouped into the “Unshared” category. All reaction norms were estimated across the entirety of each species’ range. Binned parameter estimates are available in Figs. S2–S3.

### H2 results

For species common to the two datasets, reaction norms (slopes of phenology vs. spring temperature) were generally similar between datasets when analyzed using the common framework (Fig. 2; 95% Bayesian credible intervals of slopes overlap for 4 of 6 species). However, the ASF dataset consistently showed lower temperature sensitivity of trees (Fig. 2B), and higher temperature sensitivity of wildflowers (Fig. 2A), compared to the MHKP dataset. This pattern mirrors the overall difference between the original studies (Fig. 1A) and suggests that the difference cannot be explained by the inclusion of different species in each study. A similar pattern can be seen for the species not shared between the two datasets: again, the ASF dataset shows trees to be overall less sensitive (Fig. 2B, having generally shallower reaction norms) and wildflowers to be more sensitive (Fig. 2A), compared to the MHKP dataset. Thus, again, the overall pattern does not seem to be due to species selection.

### H3. Differences in spatiotemporal extent

Shifts in phenology are dependent on environmental variation in drivers such as temperature and precipitation. While many global change studies focus on how environmental conditions vary over time (Fu et al., 2012; Kudo et al., 2004; Zohner et al., 2018), there is also substantial literature showing how geospatial variation in environmental conditions affects the timing of flowering and leaf-out both within a given growing season and in terms of long-term temperature sensitivity (Peaucelle et al., 2019; Wang et al., 2023). Moreover, several of these studies focus specifically on how temporal and spatial environmental variation interact to influence the phenology of both plants and animals (Kharouba et al., 2018; Kudo & Ida, 2013; Wann et al., 2019).

We found substantial differences between the two studies in both the spatial and temporal extent of the data, with the latter difference being more extreme due to the data sources involved. ASF collated all observations from the USA–National Phenology Network database (https://data.usanpn.org/observations/get-started; hereafter ‘NPN’) from January 1 to July 31 in the years 2009–2021, limiting their dataset to locations from latitudes 35° to 48.2° N (based on data availability) and longitudes east of 91° W (reflecting the range of eastern hardwood forests). NPN only includes observations made in the United States, so the authors could not include any Canadian observations in their analysis. In contrast, MHKP included observations dating from 1870 to 2019, with more observations in the south than in the north, ranging from 27.2° to 48.6° N. Their dataset included observations from southern Canada and from as far west as 96.4° W.

Even though the MHKP dataset includes much older records than the ASF dataset, the former includes many more datapoints representing warm spring temperatures (Fig. 3B), due to the inclusion of records from more southern locations. Conversely, despite the similar northern limits of the two studies, the ASF dataset includes more datapoints representing cool springs (Fig. 3). Thus, the two studies capture largely overlapping but distinct spatial portions of the overall spring phenology–temperature relationship for each plant functional group, with the MHKP dataset tending to skew towards warmer temperatures than the ASF dataset. If relationships between phenology and spring temperature are in fact non-linear (cf. Iler et al., 2013), but were modelled as linear by each team of researchers, the different slope estimates could simply be the result of quantification over different portions of the observed environmental variation—and would consequently be valid only over the range of the data each group considered.

**Fig. 3.**
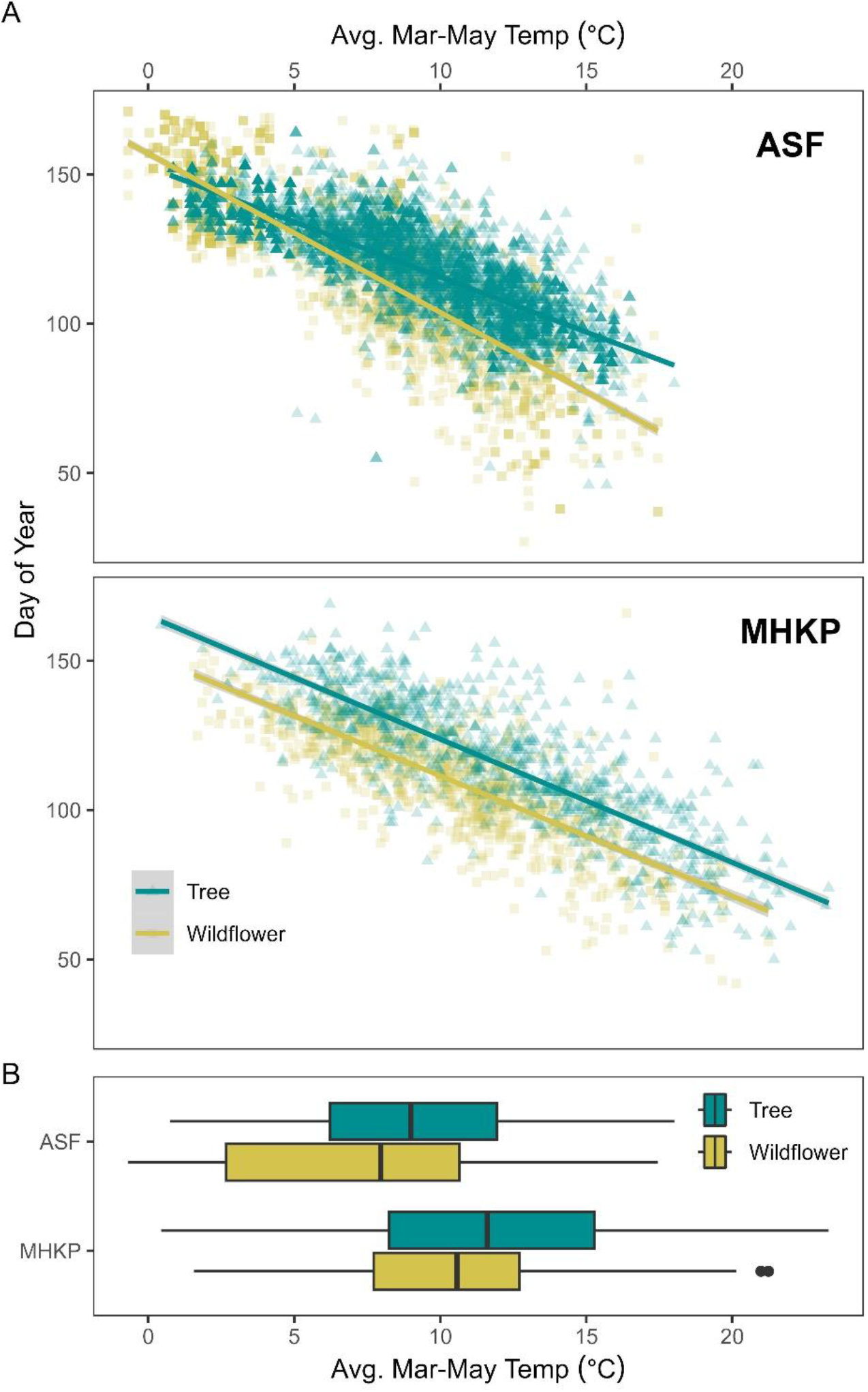
(A) Spring phenology vs. springtime (March–April) temperatures based on the Alecrim et al. (2023; “ASF”) and Miller et al. (2022; “MHKP”) datasets. Green triangles represent observations of tree leaf-out (ASF) or presence of young leaves on trees (MHKP); yellow squares represent observations of wildflower leaf-out (ASF) or presence of open flowers on herbaceous plants (MHKP). Lines are simple linear fits, with 95% confidence intervals, for each functional group; they do not account for effects of species, year, or locality. (B) Boxplots show the range (whiskers), first and third quartiles (box), mean (center line of box), and outliers (points) of estimated March–May temperatures in the Alecrim and Miller datasets. Different colored boxes differentiate between temperatures in the tree (green) and wildflower (yellow) datasets. Note the greater number of observations with temperatures >18°C in the MHKP dataset, and the greater number of observations with temperatures <5°C (particularly for wildflowers) in the ASF dataset, reflecting the difference in the latitudinal range.

To evaluate the extent to which differences in spatiotemporal extent influenced the results in these two studies, we cropped the original datasets so that data from both MHKP and ASF datasets matched in 1) spatial or 2) spatial and temporal extent. That is, for a simpler and more direct test of the impact of having different spatiotemporal extents, we re-ran the original models described in Eq. 3 using either 1) only the data from the geographic range delimited in the original ASF analysis, or 2) only the data from the same temporal *and* geographic range as the ASF dataset.

### H3 results

Cropping the MHKP dataset to match the spatial (Fig. 4A) or spatial and temporal (Fig. 4B) extent of the ASF dataset moved the estimated difference between wildflower and tree phenological sensitivity (red circles) closer to that estimated in the ASF analysis when compared to using only the hybrid model structure (Fig. 1B). (Note that because the years and the geographic extent of the ASF dataset were each a subset of those encapsulated by the MHKP dataset, the ASF dataset was not cropped for this analysis and so the parameter estimates in Fig. 4 are the same as those reported in Fig. 1B.) The estimated mean phenological sensitivity difference in the MHKP analysis moved closer to zero when data were cropped by latitude and beyond zero into negative values when cropped by both latitude and year. In addition to these shifting mean values, 95% Bayesian credible intervals also became much wider when compared to the results of the main model (Eq. 3, Fig. 1B). This is likely primarily due to substantial reductions in sample size: the MHKP dataset was reduced from n = 2016 to n = 1582 (21.5% reduction) when cropped to the same spatial extent as the ASF dataset (n = 8045, no change from cropping) and to n = 47 (97.7% reduction) when cropped to the same spatial and temporal extents as the ASF dataset. Cropping the MHKP dataset thus led to drastic changes in the model’s ability to converge during the analysis of data split into latitudinal bins, due to data in some bin × group combinations being depleted to the point where meaningful inference was impossible (Figs. S4-S5). Overall, these results suggest that differences in spatiotemporal extent of the datasets may be at least partially responsible for the differences in initially-reported model results. However, the reduction in sample size needed to run the analysis prevents us from making stronger conclusions.

**Fig. 4.**
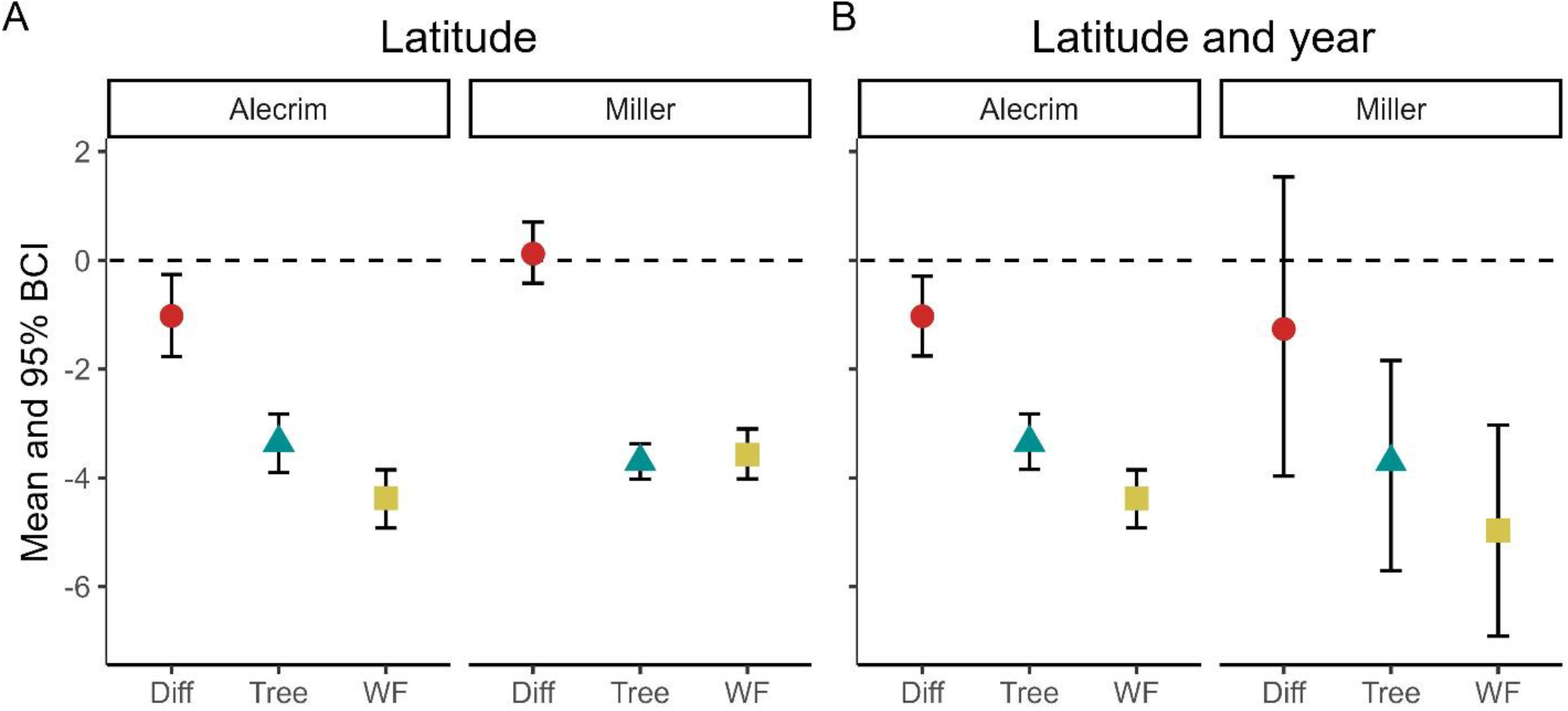
Posterior estimated means (points) and 95% Bayesian credible intervals (whiskers) of phenological sensitivities of wildflowers (WF; yellow squares), trees (green triangles), and the difference between them (red circles) using the ASF and MHKP datasets. Panel A shows posterior estimates for when the datasets are cropped to the same geographic extent and panel B shows estimates for when datasets are cropped to the same spatial and temporal extents. The extents of the ASF dataset were subsets of the extents of the MHKP dataset and so were not cropped. Posterior parameter estimates for latitudinal/temperature bins for versions of the cropped datasets are provided in Fig. S4-S5.

### H4. Differences in phenophase

Previous studies have shown that different phenophases have different temperature sensitivities (Buonaiuto et al., 2021; Geng et al., 2022). The MHKP and ASF studies differed regarding the phenophases used, which could also contribute to differences in the results (Table 1). Specifically, while both original studies used some measure of canopy tree leaf-out phenology, the MHKP study measured phenological sensitivity of wildflower flowering whereas the ASF study measured changes in wildflower leaf-out. To test whether these differences may have resulted in the different results, we ran the ASF models (following the model structure in Eq. 3) but with the phenophase described by NPN as “open flowers” (Denny et al., 2014) for wildflowers, keeping the phenophase “leaf-out” for trees. It should be noted that data for the “open flowers” phenophase are available for only a 10-year timespan (vs. the 13 years of leaf-out data used by ASF). We did not change the phenophases used in the MHKP dataset.

### H4 results

Using this approach, neither the ASF nor MHKP datasets showed a statistically significant difference in phenological response between wildflowers and trees (Fig. 5; 95% Bayesian credible intervals overlapped zero). However, the ASF dataset still indicates a trend toward greater thermal sensitivity in wildflowers (negative values), while the MHKP dataset still indicates the opposite trend.

**Fig. 5.**
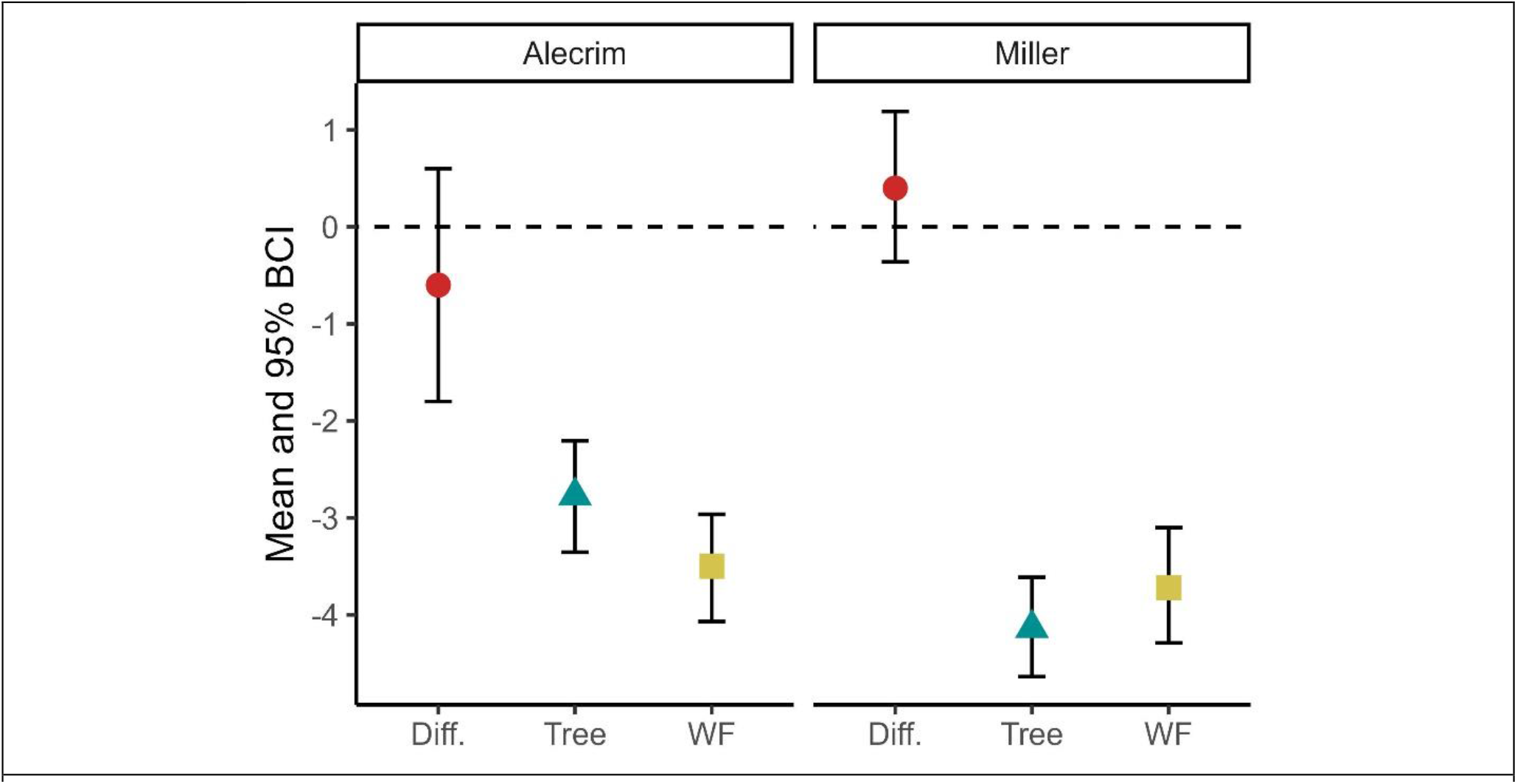
Posterior estimated means (points) and 95% Bayesian credible intervals (whiskers) of phenological sensitivities of wildflowers (WF; yellow squares), trees (green triangles), and the difference between them (red circles) for the ASF (left panel) and MHKP (right panel) datasets. The ASF posteriors are from the model using the flowering (“open flowers”) NPN phenophase, whereas the MHKP posteriors are the same as presented in Fig. 1B for the ease of comparison.

## Conclusions

We present a series of analyses to test a set of defined hypotheses with the aim of reconciling the contradictory results from two recent studies concerning the sensitivity of canopy tree phenology and understory herb phenology to warmer spring temperatures. One study (Miller et al., 2022) originally found that trees are more sensitive to warming than herbs, whereas the second study (Alecrim et al., 2023) found that herbs are more sensitive than trees. We hypothesized that one or some combination of four different hypotheses may be responsible for these discrepancies, particularly given recent evidence suggesting strong agreement in phenological signals between herbarium and other datasets (Iwanycki Ahlstrand et al., 2022; Park et al., 2019; Ramirez-Parada et al., 2022).

Overall, we find that model structure appears to be a major factor in the different conclusions reached by the two studies. Specifically, the Miller et al. result, that trees are more responsive to temperature change than herbs, is no longer supported once the random slope estimates are included to match the model structure of the Alecrim et al. approach (Fig. 1B). Merging the model structures also moved the reaction-norm difference described in Alecrim et al. closer to zero, although it remained significantly negative (indicating higher sensitivity in spring wildflowers; Fig. 1B). Thus, although the hybrid model structure brought the two results closer together, there remained discrepancies that were not explainable by model structure, environmental data source, or parameterization of spring temperature windows alone.

Differences in the thermal range extent of the two datasets also seemed to explain some of the contradicting results: if the datasets are both constrained to only include locations used in Alecrim et al., the results converge (Fig. 4). However, the caveat to this finding is that the Alecrim dataset only included a subset of the spatial and temporal extent of the Miller dataset, meaning that only the Miller dataset was cropped for this part of the analysis. Although the mean response in the Miller dataset converged toward the Alecrim result (i.e., a signal of wildflowers being more sensitive to spring temperature compared to trees), the sample size was greatly reduced and as a result 95% Bayesian credible intervals greatly widened. Therefore, while we conclude that this hypothesis is supported to some extent, we note that convergence could be due instead to a lack of good herbarium data over the 12-year period represented in the Alecrim dataset.

Differences in species selection did not seem to explain much, if any, of the difference in the results, suggesting that group-level signals are relatively strong (at least within each dataset, contingent on the differences already described). Species that were shared across both analyses were generally found to have similar sensitivities to the March–May temperature window (Fig. 2). Although this only included six of the total 28 species investigated across both analyses, the species unique to each study also did not appear to strongly differ in their overall sensitivities (‘Unshared’ species in Fig. 2), suggesting that results would not have been strongly affected in either study had a different group of species been selected for analysis.

Furthermore, when controlling for model structure, temperature window, and choice of phenophase, the differences between the two studies still could not be explained. The ASF dataset still points towards an overall trend of wildflowers being more sensitive to temperature, while the MHKP dataset points towards the opposite trend, i.e., trees being more sensitive. Again, model structure played an important role in determining the results. This is especially true for the MHKP dataset, where the difference between the phenological response between wildflowers and trees is detectable only when using the original model structure (Fig. 1A), but not after merging the model structures (Fig. 1B).

The main difference between model structures was the inclusion of random slopes. Random slopes are typically used when analyzing data that involves repeated measurements over time or across different groups. This approach allows the estimation of individual-level variability in the relationship between the temperature (the predictor variable) and leaf-out/flowering day (outcome). They are also helpful when analyzing data with a hierarchical structure, such as when measurements are taken within nested groups (e.g., species within a functional group). In these cases, random slopes can account for within-group variation and improve the accuracy of the analysis (Oberauer, 2022). For the ASF study, a random slope for the interaction between species and site was included to account for the variability in the response to temperature caused by differences between species and geographical locations. The decision from the authors of the MHKP structure to use nonhierarchical, shared slopes was based largely on precedence from previous models of phenological sensitivity that used this approach (e.g., Heberling et al., 2019). This approach did not allow Miller et al. (2022) to evaluate species-level sensitivities, but instead places the model focus on community signals (grouping the data from all species together). Overall, the decision to use random slopes in Bayesian analysis should be based on the specific characteristics of the data and research question at hand.

Importantly, although we focus here on overall results (i.e., the posterior estimated parameter values when considering the entirety of each dataset), we also want to return to the initial goal of these studies, which was to investigate whether phenological sensitivity of wildflowers and trees varied across the extent of North America. In their initial studies, both sets of authors found that phenological sensitivities, and therefore phenological escape dynamics, differed according to latitude or regional climate. Alecrim et al. (2023) found no difference in sensitivity between wildflowers and trees in the southern part of the range investigated but found wildflowers to be relatively more sensitive in the central and northern regions (Fig. S1A). Miller et al. (2022) also found no difference in sensitivity between wildflowers and trees in the southern part of the range but found no significant difference between tree and wildflower sensitivity in the central and northern portions of the range either (Fig. S1A). A unique goal of Miller et al. (2022) was also to explore differences with the shrub layer and between native and non-native species.

Upon merging the models into the hybrid model structure (Eq. 3), these binned responses changed in unexpected ways. While the Miller et al. binned results did not change with respect to overall signal (trees were more sensitive than wildflowers in the southern bin, with no significant difference between functional groups in the central and northern bins), the Alecrim et al. results changed such that differences in sensitivity between wildflowers and trees were no longer significantly different in any of the bins (Fig. S1B). We are thus left with seemingly contradictory results, wherein the overall signal in the Miller paper was more sensitive to changes in bin structure (e.g., cropped vs. uncropped; binned by latitude vs. binned by spring temperature), while the binned results themselves seemed to be relatively robust. The opposite was true for the Alecrim dataset. Therefore, while the changes in the total dataset results appear to support the notion that the Alecrim results are more robust with respect to trends estimated across the entire range of the study area (eastern North America), the Miller results were more robust at the latitudinal/ temperature bin level.

Where does this leave us with reconciling the contradictory results in these two models? Starting with where the datasets seem to agree, both seem to consistently indicate that wildflowers in the southern/warmer portion of the geographic ranges present in the datasets (eastern North American temperate deciduous forest) are more likely to have lower relative phenological sensitivity than co-occurring tree species. Furthermore, this result appears to be relatively robust regardless of which medley of species are included in the analysis. Importantly, both datasets suggest strikingly similar estimates of the sensitivity of wildflowers and trees to spring temperature (95% Bayesian credible intervals largely overlap among species within groups and between the two studies). Further, phenological activity was consistently found to be earlier in warmer years and warmer locations in a quantitatively similar way using both community science and herbarium datasets. This is largely consistent with previous studies that have found similar parallels when comparing herbarium and community-generated observational datasets (Iwanycki Ahlstrand et al., 2022; Ramirez-Parada et al., 2022; Spellman & Mulder, 2016).

However, full reconciliation of model differences remains elusive. While merging model structure, environmental data source, and binning approach all tended to move the results closer together, the datasets still produced results that suggested different ecological trends. Specifically, the Alecrim et al. (2023) dataset consistently suggests that wildflowers are generally more sensitive to spring temperature compared to trees and the Miller et al. (2022) dataset suggests that wildflowers are either as sensitive as or less sensitive than tree species. Cropping the Miller dataset to match the same spatiotemporal extent as the Alecrim dataset did appear to fully reconcile the model differences (Fig. 4), but the reduction in sample size led to a sharp increase in model uncertainty (including in the binned model; Figs. S4-S5) and so we hesitate to make firm assertions based on that analysis. Moreover, the reduced sample size prevented some of the binned models from converging when using the cropped datasets (i.e., the Cool bin in the Miller dataset; Fig. S5), preventing us from making any inference about this aspect of model performance. Lastly, cropping the MHKP dataset also disproportionately removed data from warmer (more southern) regions, potentially leading to new biases associated with reductions in predictability under warmer climate conditions.

In sum, our analyses demonstrate that choices of model structure can greatly impact study conclusions, a point that has been raised previously in ecology (e.g., Arnqvist, 2020). In addition, although our results seem to support recent findings that community science data and herbarium collection data yield analogous estimates of phenological sensitivity in North America at the species level (Iwanycki Ahlstrand et al., 2022; Ramirez-Parada et al., 2022; Spellman & Mulder, 2016), they also suggest that inferences about complex ecological mechanisms such as species interactions may still be significantly affected by the type and spatiotemporal extent of data. Still, until crowd-sourced data repositories grow to match (or at least approach) the climatic coverage present in herbarium collections or until herbarium collections grow to capture the same spatial extent and temporal frequency of community science repositories, we conclude that ecological studies should incorporate both datasets in analysis as much as possible to rule out the possibility that one or the other biases inference.

## Author contributions

BRL and EFA led the statistical analysis. All authors contributed equally to the writing and interpretation of results.

## Data availability

Datasets used in this analysis are available in their entirety with the original manuscripts we compare here.

## Acknowledgements

We thank our Tara Miller and Sara Kuebbing, coauthors of the original research articles compared in this article. We thank the PRISM climate group and the National Phenology Network for making their data freely available for use in this study. This work was supported by the Ontario Ministry of Research, Innovation and Science through an Early Researcher Award to JF, a Natural Sciences and Engineering Research Council of Canada (NSERC) Discovery Grant to RS, and National Science Foundation Grants No. DEB 1936971 to JMH and 1936877 to RBP and supporting BRL.

