## Supporting information for "Phenological mismatch between trees and wildflowers: Reconciling divergent findings in two recent analyses"

**Fig. S1:** Posterior means (points) and 95% Bayesian credible intervals (whiskers) for the estimated difference in phenological sensitivity (red circles) between wildflower (yellow squares) and canopy tree (blue triangles) sensitivity in A) the original models and B) the hybrid model structure described in Eq. 3. Filled points represent the posterior estimated means for the NPN data used by Alecrim et al. (2023) whereas open points represent those for the herbarium dataset used by Miller et al. 2022. Different binning approaches were used in the original models (described in Table 1) and the hybrid model structure used a latitudinal binning approach described in the Methods section. Warm and Cool bins correspond to low- and high-latitude bins used by Alecrim et al. (2023), respectfully.

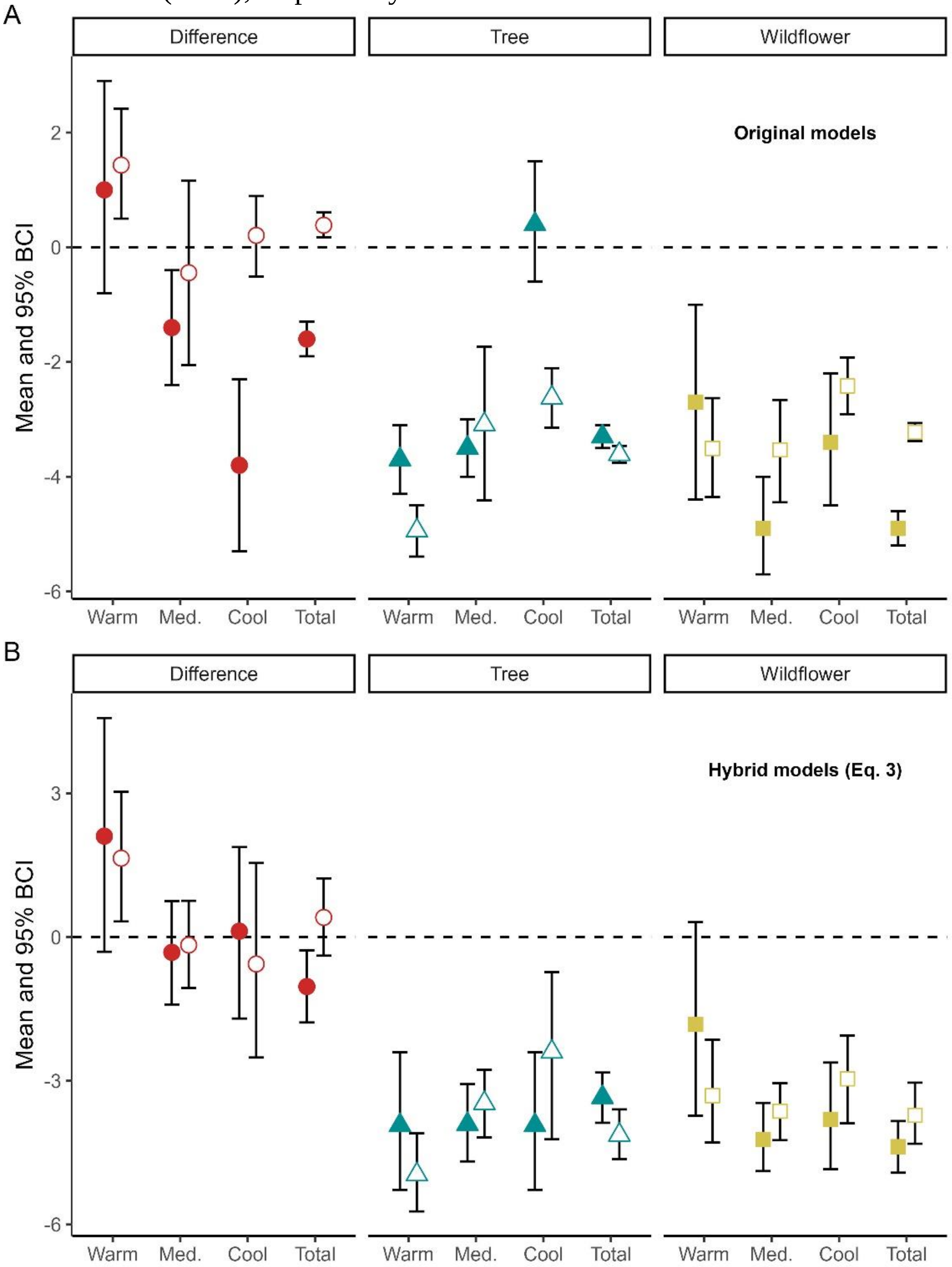

**<u>Fig. S2</u>**: Binned posterior estimates of the mean (points) and 95% Bayesian Credible Intervals (BCI, whiskers) phenological sensitivity for the two wildflower (top row) and four canopy tree species (bottom two rows) shared between Alecrim et al. (2023, red circles) and Miller et al. (2022, blue triangles). Posterior estimates were calculated using the hybrid model structure (Eq. 3). Warm and Cool bins correspond respectfully to the low- and high-latitude bins used in the hybrid model structure.

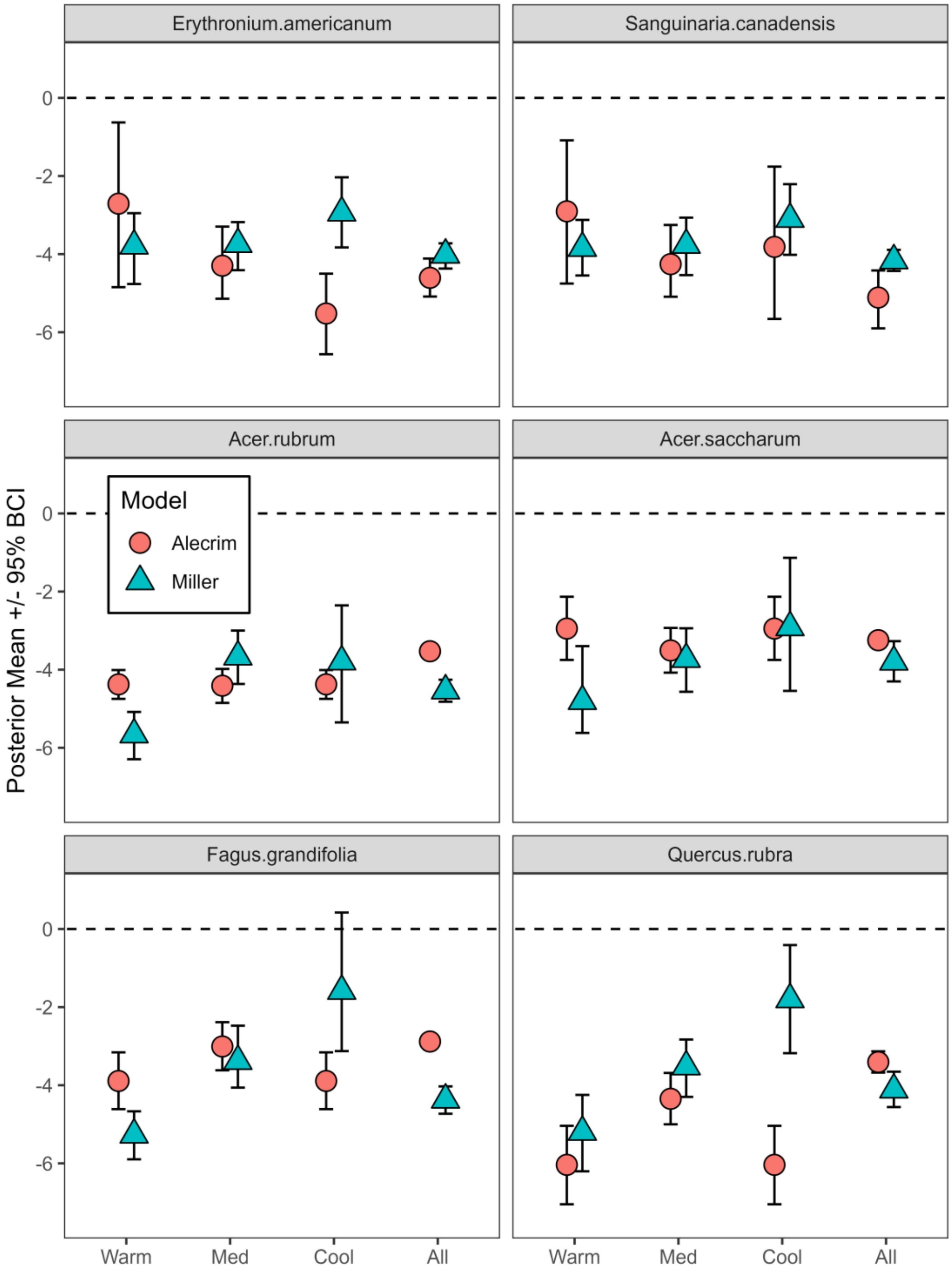

**<u>Fig. S3</u>**: Binned posterior estimates of the mean (points) and 95% Bayesian Credible Intervals (BCI, whiskers) phenological sensitivity for the 13 wildflower (top panel) and eight canopy tree species (bottom panel) shared between Alecrim et al. (2023, red circles) and Miller et al. (2022, blue triangles). Posterior estimates were calculated using the hybrid model structure (Eq. 3). Warm and Cool bins correspond respectfully to the low- and high-latitude bins used in the hybrid model structure.

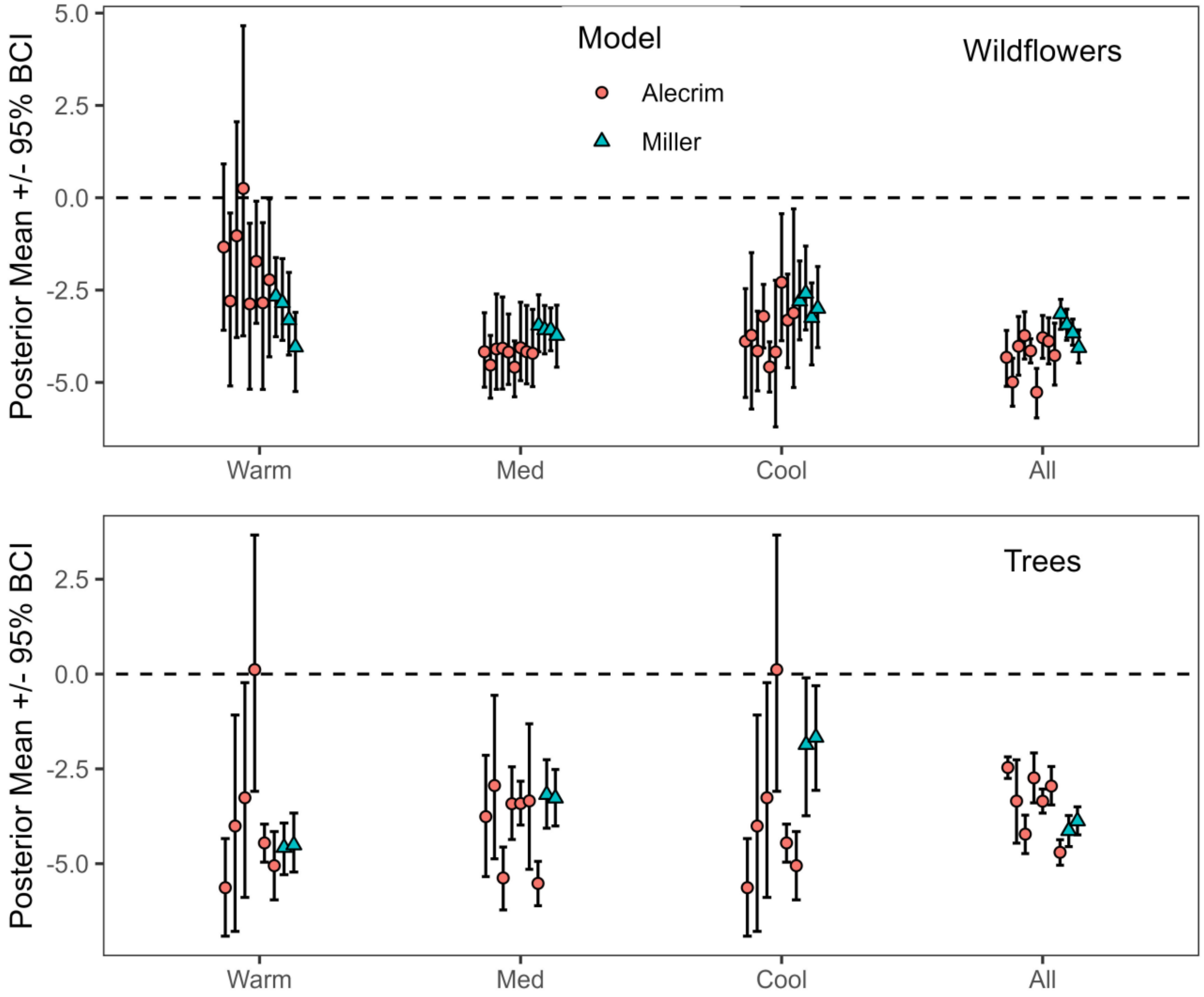

**Fig. S4:** Posterior means (points) and 95% Bayesian credible intervals (whiskers) for the estimated difference in phenological sensitivity (red circles) between wildflower (yellow squares) and canopy tree (blue triangles) sensitivity when the Miller et al. (2022) dataset (empty points) is cropped to the same spatial extent as the Alecrim et al. (2023) dataset (filled points). Model structure follows Eq. 3. Warm and Cool bins correspond to low- and high-latitude bins used by Alecrim et al. (2023), respectfully.

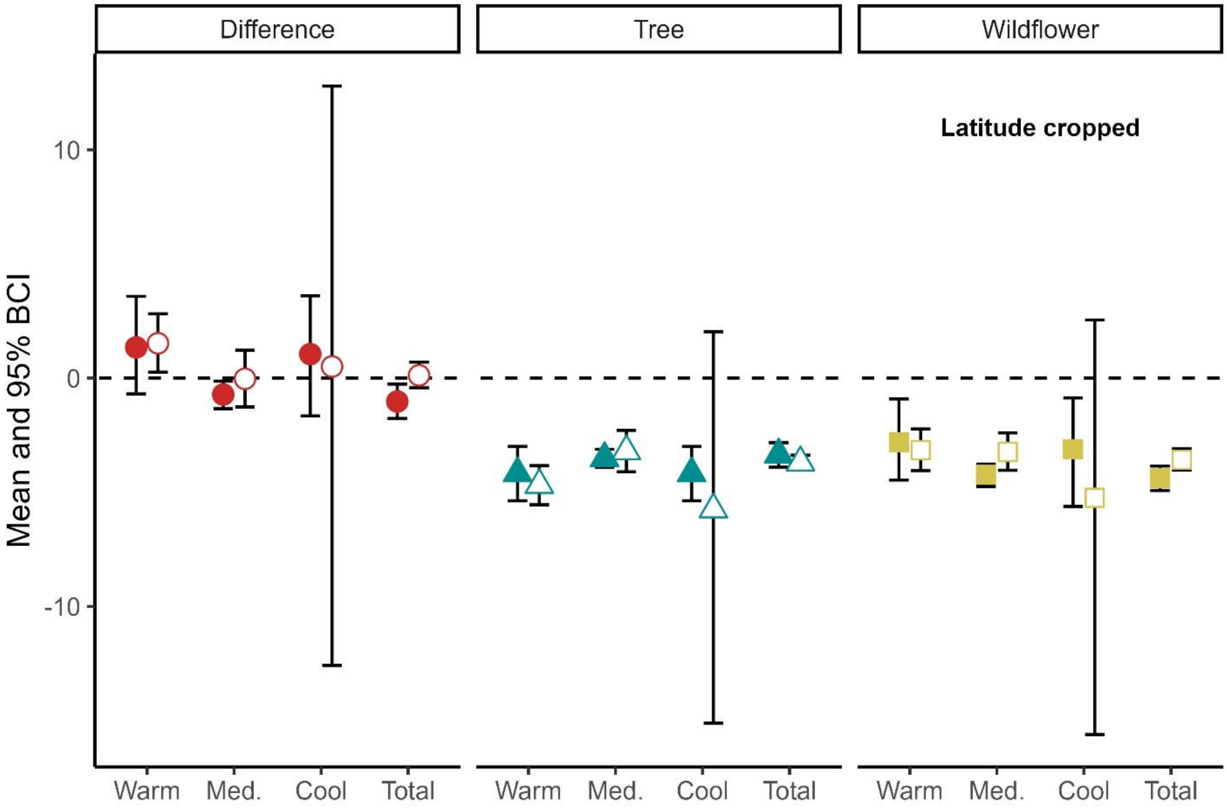

**<u>Fig. S5:</u>** Posterior means (points) and 95% Bayesian credible intervals (whiskers) for the estimated difference in phenological sensitivity (red circles) between wildflower (yellow squares) and canopy tree (blue triangles) sensitivity when the Miller et al. (2022) dataset (empty points) is cropped to the same spatial and temporal extent as the Alecrim et al. (2023) dataset (filled points). Model structure follows Eq. 3. Warm and Cool bins correspond to low- and high-latitude bins used by Alecrim et al. (2023), respectfully. Values for the Miller et al. (2022) dataset Cool bin are missing because the model did not converge.

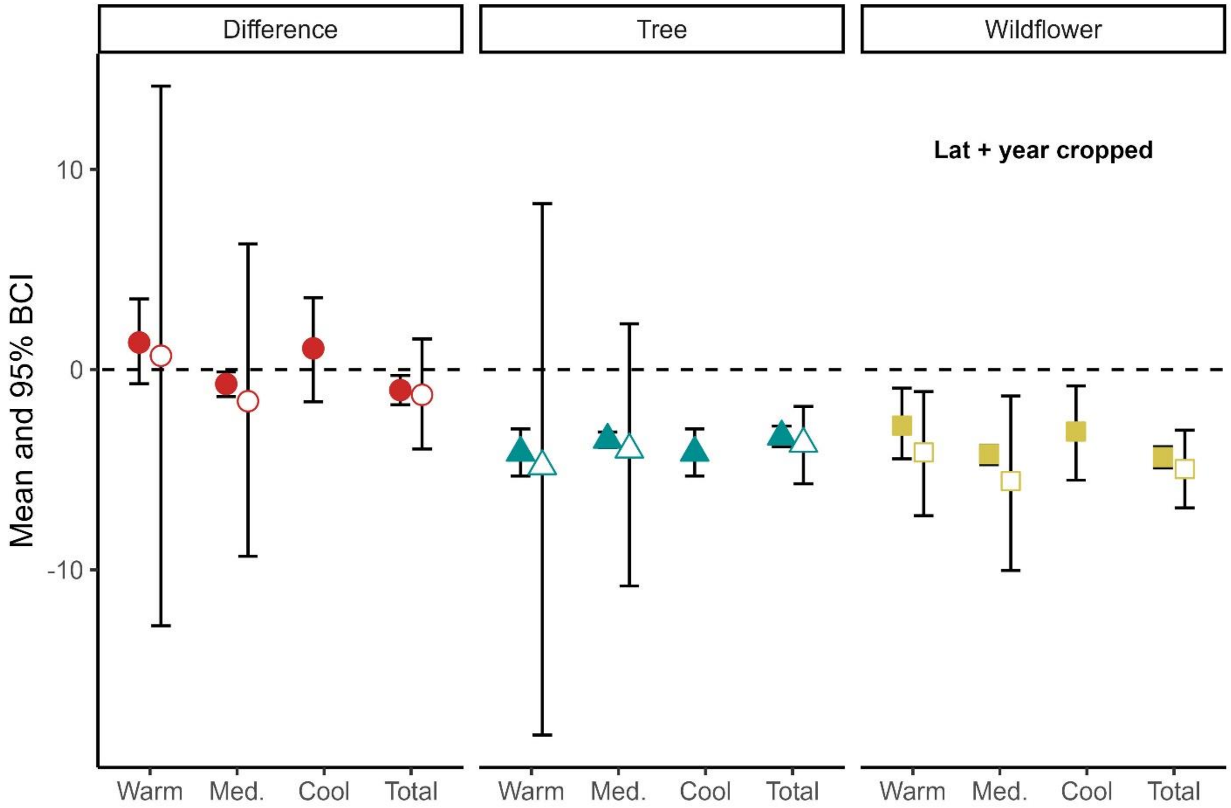
